# The Bacterial Replicative Helicase Loader DciA is a DNA Condenser

**DOI:** 10.1101/2023.09.08.556801

**Authors:** Stéphanie Marsin, Sylvain Jeannin, Sonia Baconnais, Hélène Walbott, Gérard Pehau-Arnaudet, Magali Noiray, Magali Aumont-Nicaise, Emil GP Stender, Claire Cargemel, Romain Le Bars, Eric Le Cam, Sophie Quevillon-Cheruel

**Author notes:** New address: Centre de Biologie Structurale (CBS), Université Montpellier, CNRS, INSERM, Montpellier, France.

## Abstract

The loading of the bacterial replicative helicase is an essential step for genome replication and depends on the assistance of accessory proteins. Several of these proteins have been identified across the bacterial phyla. DciA is the most common loading protein in bacteria, yet the one whose mechanism is the least understood. We have previously shown that *Vc*DciA from *Vibrio cholerae,* composed of a globular KH-like domain followed by an unfolded extension, has a strong affinity for DNA. Here, we characterized the droplets formed by *Vc*DciA upon interaction with a short single-stranded substrate. We demonstrate the fluidity of these droplets using light microscopy and address their network organization through electron microscopy, thereby bridging events to conclude on a liquid-liquid phase separation behavior. Additionally, we observe the recruitment of *Vc*DnaB inside the *Vc*DciA-DNA droplets. We show that DnaC from *Escherichia coli* is also competent to form these condensate structures in the presence of ssDNA. Our data open up possibilities for the involvement of DciA in the formation of non-membrane compartments within the bacterium, facilitating the assembly of replication players with the chromosomal DNA.

## Introduction

Replicative Helicases are essential proteins implicated in DNA replication in all kingdoms of life. They unwind the DNA double helix in front of the replisome, a large multi-subunit complex in charge of DNA strand synthesis ^1^, allowing its progression at the replication fork during the chromosome replication. Replisome assembly is tightly regulated, and the recruitment and loading of the replicative helicases on DNA is a key step in this mechanism ^2,3^.

In bacteria, the loading of the closed and circular hexameric helicase DnaB depends on the intervention of loading proteins. In *Escherichia coli* (*Ec*), DnaC, a member of the ATPase Associated with various cellular Activities (AAA+) superfamily, interacts with the LH/DH module of the C-terminal ring of DnaB as a hexamer. This interaction causes a torsion to the closed ring formed by DnaB, cracking it open into a loading-competent state enabling the loading of the helicase on *oriC* ^4–7^. In *Bacillus subtilis*, the other bacterial model studied closely, DnaI, a homolog of DnaC, has been described as a ring-maker but the molecular mechanism remains to be determined ^8^. However, DnaC and DnaI are poorly represented in the bacterial world. Indeed, it has been shown in 2016 that the genes encoding these proteins were acquired by horizontal transfer from phage genomes ^9,10^ and that the ancestral gene, replaced in these phyla by DnaC/I, encodes the protein DciA. Using DciA and DnaB from *Vibrio cholerae* (*Vc*) as models, and by comparison with the *E. coli* system, we previously demonstrated by biochemistry experiments supplemented by a structural study that DciA is indeed the helicase loader in this organism ^11^.

*Vc*DciA is a small protein of 18 kDa containing a high level of arginine and lysine residues (13 R and 13 K over 158 residues), which gives it an isoelectric point of around 10. Structural analysis by NMR revealed that the N-terminal domain of *Vc*DciA is folded as a KH fold ^11^, described in the literature to interact with nucleic acids ^12^. *Vc*DciA can indeed interact with single and double-stranded DNA, with a preference for the junction between these two structures, as we have previously shown by NMR and electron microscopy ^13^. The C-terminal domain, for its part, is unfolded ^14^, but structures into 2 small α-helices upon contact with the LH/DH module of DnaB, without applying however the torsion force observed for *Ec*DnaC on *Ec*DnaB ^15^. Moreover, we observed that in comparison with the *E. coli* system, *Vc*DciA is rapidly released from the helicase-DNA complex once the helicase is loaded on DNA.

The mechanism of DnaB loading by DciA and the different steps to reach this point remain to be deciphered. As the structure of the *Vc*DnaB•*Vc*DciA complex was restricting for the description of loading-competent state and as there is no evidence for the ability of DciA to open the DnaB ring by itself, we consider the possibility of involving a third partner in the loading mechanism of the helicase. Knowing the affinity that DciA has for DNA, we investigate how this molecule could be directly involved in the mechanism other than as the receptacle. Over the past decade, numerous studies revealed the implication of membrane-less compartments in essential processes in cells. These condensates are mostly composed of DNA and proteins, and there is an increasing interest in comprehending the role of biological condensates in these processes. The majority of observations have been made in eukaryotic cells, including the processes of DNA replication or DNA modulation by the topoisomerase ^16,17^ among many other cellular functions. A growing number of studies also report a coordination of DNA Damage Response (DDR) with Liquid-Liquid Phase Separation (LLPS) processes ^18^. A recent study described the phase separation properties of replication protein A (RPA), whose described function to protect ssDNA is now extended to genome organization and stability ^19^. Numerous studies on viruses have also been published, revealing the importance of membrane-less replication compartments ^20^. Some studies have been made in prokaryotes, where the idea of non-membrane compartmentalization allowing the concentration of proteins in specific sites of the cell via LLPS processes is advanced in different processes as varied as CO2 concentration, transcription control, DNA segregation, RNA metabolism, or cell growth and division ^21^. With respect to DNA replication, repair, and recombination, it has been shown that the single-strand DNA binding protein SSB, the bacterial RPA analog, is able to produce phase separation via protein-protein interactions ^22,23^.

In order to advance our understanding of the bacterial replicative helicase loading mechanism by the ancestral loader DciA, we investigate if DciA from *V. cholerae*, due to its appropriate biochemical properties, i.e. an intrinsically disordered region (IDR) in the C-terminal domain following a basic DNA-binding domain, was able to form LLPS in the presence of DNA, and if this organization could be involved/play a role in the recruitment of the DnaB helicase.

## Results and discussion

### *Vc*DciA forms LLPS in the presence of single-stranded DNA

The first evidence that *Vc*DciA had a tendency to form droplets was the results of crystallization trials performed with the purified protein. We have never been able to obtain *Vc*DciA crystals even at the highest protein concentration tested of 200 mg/mL (11 mM). In addition, a larger-than-average number of crystallization drops showing phase separations were observed and appeared to be highly spherical structures. Secondly, during our previous studies, we often observed turbidity when *Vc*DciA was mixed with a single-stranded DNA (ssDNA).

We then explored these properties and established the optimal conditions to observe and study *Vc*DciA droplets. In MES buffer at pH 6, with a NaCl physiological concentration of 150 mM, mixing *Vc*DciA with an oligonucleotide (at 10 µM) leads to a solution with high turbidity, due to the presence of droplets (Fig. 1A and Supplementary Fig. 1A). At 300 mM of NaCl, the turbidity decreases drastically, suggesting that electrostatic forces are involved in LLPS assembly (Supplementary Fig. 1A). No crowding agent like polyethylene glycol is required, and 10 µM of the protein is enough to observe LLPS. We then validated the presence of *Vc*DciA and ssDNA in the droplets by using labeled molecules. Co-localization of *Vc*DciA, labeled with ATTO 550, and a 50-nucleotide long oligonucleotide labeled with Cy5 (ssDNA- Cy5) was observed, confirming their presence in the droplets (Fig. 1B). On pluronic pre-treated glass, we observed that droplets are not always spherical and spread out on the glass (Fig. 1B and Supplementary Fig. 1B). We tested then a silane coating to obtain a neutral surface, and observed that the droplets are indeed perfectly spherical even in the vertical axis, and do not wet the glass surface under these conditions (Fig. 1C and Supplementary Fig. 1C). We observed that ssDNA-Cy5 added to preformed LLPS composed of DciA-ATTO 550 and unlabeled ssDNA quickly penetrates the droplets (Fig. 1C). The fluorescent ssDNA can be detected 10 seconds after its addition at the periphery of the droplets, and is totally integrated after 20 seconds, presenting same homogeneous distribution inside the droplets as labeled DciA and illustrating its fast diffusion.

**Figure 1:**
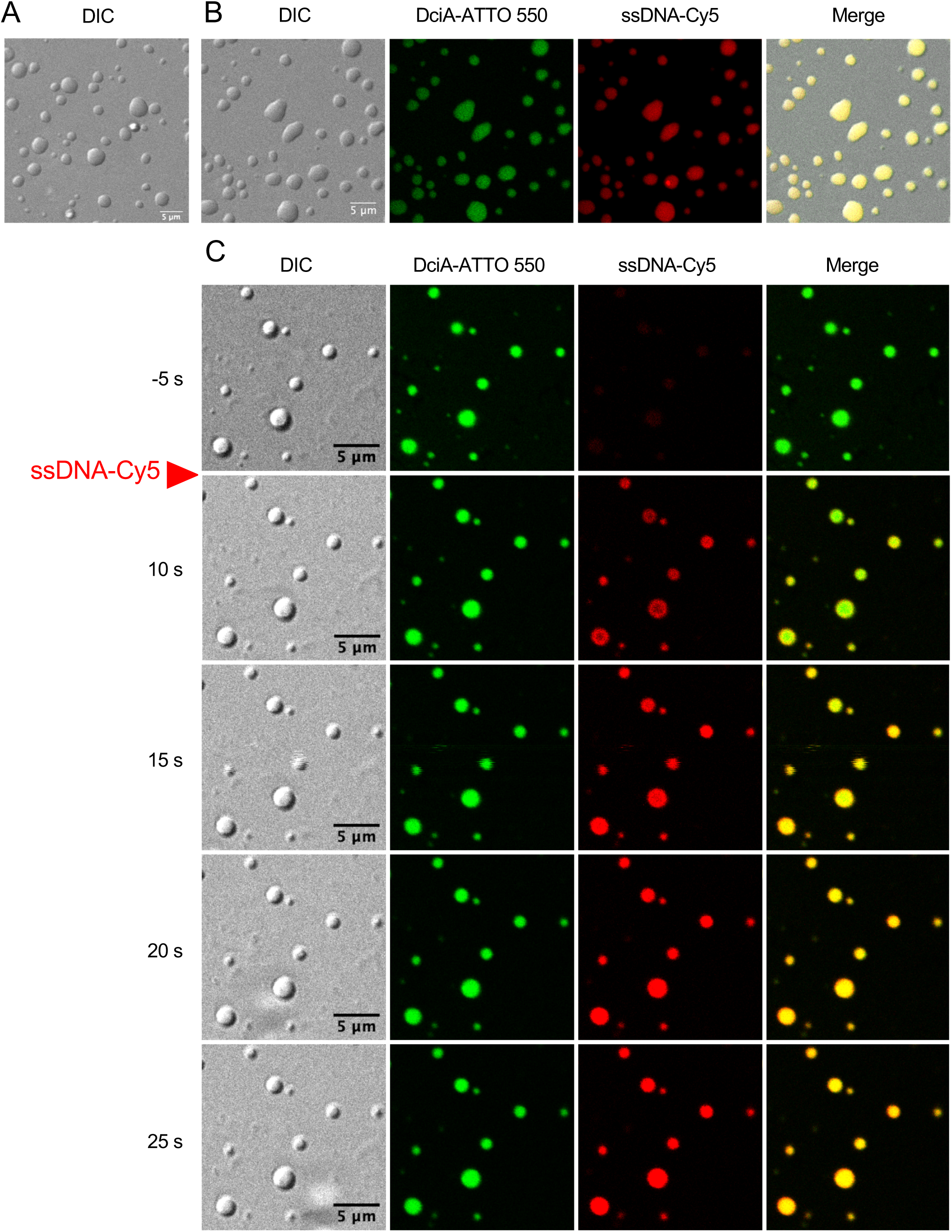
Photonic Microscopy images of droplets formed by *Vc*DciA and ssDNA. **A.** DIC images of *Vc*DciA in the presence of ssDNA. LLPS were formed with 15 µM of *Vc*DciA in the presence of 10 µM of poly- dT50 and loaded on glasses treated with pluronic alone (see Methods). 5 µM scale is indicated. **B.** Representative confocal images of *Vc*DciA labeled in the presence of fluorescent ssDNA. LLPS were formed with 15 µM of *Vc*DciA labeled with ATTO 550, in the presence of 10 µM of poly-dT50 and 0.2 µM of ssDNA labeled with Cy5 (ssDNA-Cy5). LLPS were loaded on glass treated with pluronic alone (see Materials and methods). 5 µM scale is indicated. **C.** Microscope visualization of ssDNA integration into the droplets. LLPS were formed with 15 µM of *Vc*DciA labeled with ATTO 550, in the presence of 10 µM of poly-dT50, and 0.2 µM of ssDNA-Cy5 was added at the time indicated with the red arrow. Glasses were pretreated with Sigmacote and pluronic (see Materials and methods). An image is shown every 5 seconds. A 5 µm scale is indicated.

ssDNA and *Vc*DciA are both required for droplets formation. This result is similar to that observed with ssDNA binding protein RPA ^19^ and LLPS behavior had then to be confirmed.

### Microscopy analysis of the *Vc*DciA LLPS reveals liquid behavior

The first characteristic of LLPS is their ability to fuse. We observed, under the microscope, many fusion events between droplets (Supplementary Fig. 1D). The fusions are very rapid and occur in less than 5 seconds (time between two consecutive images). We then tried to follow the fusion of two droplets labeled by different fluorescent tags: ATTO 550 and Cy5 (Fig. 2A). First, we induced droplets formation with *Vc*DciA labeled by ATTO 550 and unlabeled ssDNA (Fig. 2A, -5 s). Then we add preformed droplets containing unlabeled *Vc*DciA and ssDNA-Cy5 on the glass slide (Fig. 2A). At 133 seconds, we can see the early steps of the fusion event between an ATTO 550 droplet and a preformed Cy5 one (Fig. 2A in the circle). The size of the droplets increases, as shown in the dotted circle in the DIC channel (Fig. 2A), but the two colors remain distinct. 5 seconds later (time = 138 s), the two droplets co-localize and the fusion is achieved. Colors merge totally in the next image, 5 seconds later. On the same field of view, at 174 seconds, two smallest droplets are present (in the dotted rectangle Fig. 2A). One was preformed and only labeled by ATTO 550, while the left one, coming from an out-of-focus plane, is already labeled by both ATTO 550 and Cy5, probably due to a previous fusion event. The two droplets fuse together and relax into one single round/spherical droplet. These observations confirm the liquid-like behavior of *Vc*DciA droplet. It was also possible to observe fusion events between two proximal droplets already installed on the glass (Supplementary Fig. 1D), attesting to the versatility behavior of these structures.

**Figure 2.**
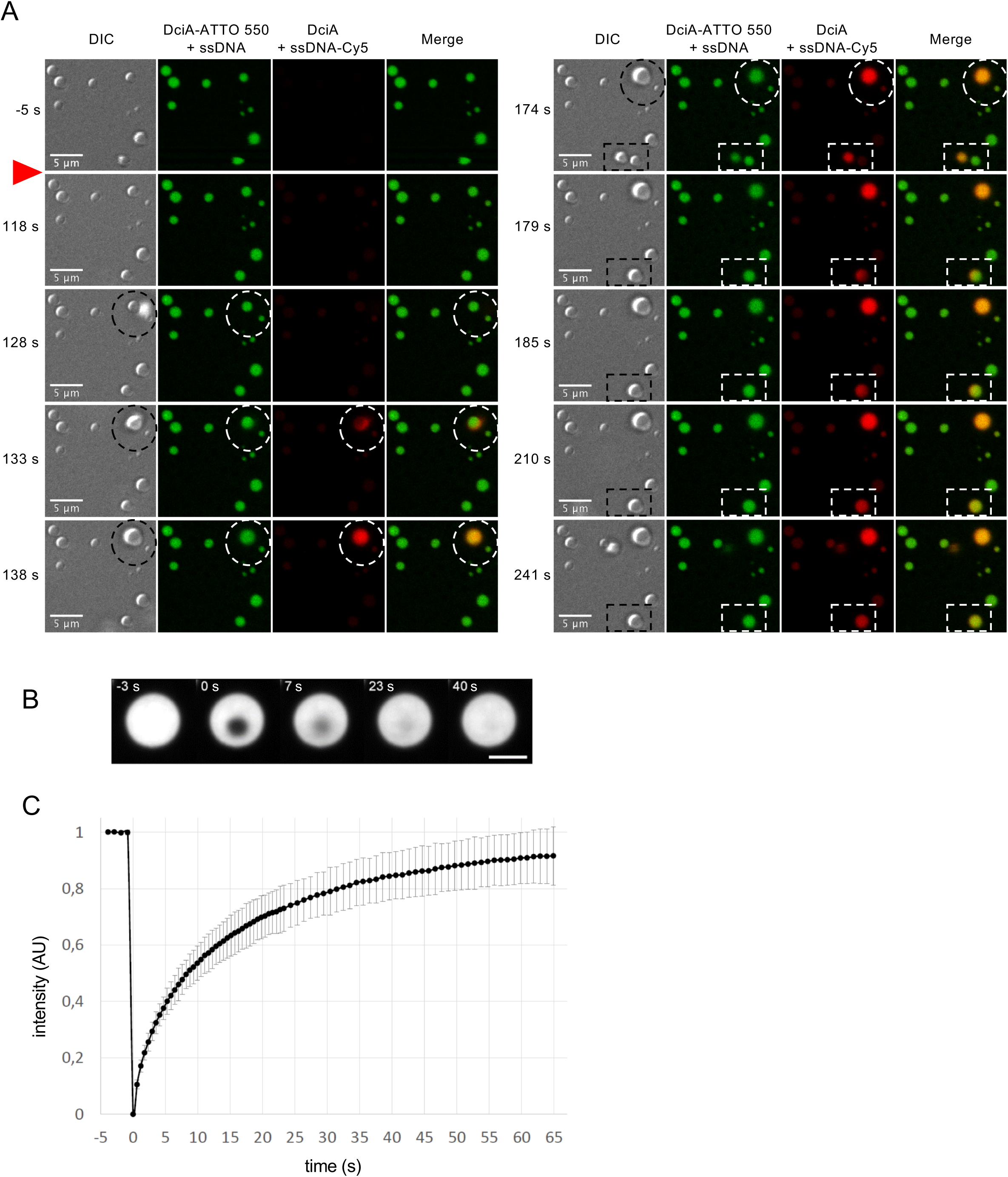
Analysis of *Vc*DciA droplets behavior by microscopy. **A.** Fusion of *Vc*DciA droplets over time. LLPS were first formed with 30 µM of *Vc*DciA-ATTO 550 in the presence of 10 µM of poly-dT50 (-5 s). Droplets preformed by *Vc*DciA with 10 µM of poly-dT50 and 0.2 µM of ssDNA-Cy5 were then added at the time indicated by the red arrow. Images collected at indicated times were presented as they illustrated fusions between droplets of different colors. Dashed circles and rectangles indicate the fusion events of droplets. A 5 µm scale is indicated. **B.** Representative time-lapse imaging of a FRAP experiment. LLPS were first formed with 30 µM of *Vc*DciA-ATTO 550 in the presence of 10 µM of poly- dT50. FRAP was applied on a central sub region (1 μm diameter). Time is indicated in seconds. The bleaching event is performed at time 0s. The scale bar is 3 μm. **C.** Quantification of recovery during FRAP experiments. Representation of the mean fluorescence intensity recovery of FRAP experiments presented in B (±Standard deviation). Data were obtained from 3 independent replicates (n=34).

To further demonstrate the liquid-like dynamics of the *Vc*DciA droplets, we performed Fluorescence Recovery After Photobleaching (FRAP) experiments on *Vc*DciA labeled with ATTO 550 (Fig. 2B). *Vc*DciA exhibits fast dynamics with a nearly complete recovery after 1 minute of imaging: the mobile fraction represents 97.30±11%, with a half-time recovery 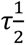= 6.81±1.9 seconds for a bleach spot of 1 μm (Fig. 2C). Such a diffusion rate is consistent with those of other membrane-less organelles, such as nuclear speckles and nucleoli ^24,25^.

For a better understanding of the electrostatic interactions inside the droplets, we performed FRAP experiments on both the ssDNA-Cy5 and *Vc*DciA-ATTO 550 at the same time. The ssDNA-Cy5 also showed a fast recovery, however, this recovery was slightly slower 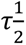= 49.88±20 s and less complete than *Vc*DciA (only 76.18±16% after three minutes) (Supplementary Fig. 2A and 2B). If we compare the 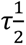 for each bleaching event, we measured that the DNA mobility is 3.61 fold slower than those of the *Vc*DciA protein (Supplementary

Fig. 2C). The fact that the DNA mobility is slower than *Vc*DciA and incomplete after three minutes, suggests that additional interactions, in addition to electrostatic forces, limits its dynamic inside the droplets. This is consistent with the hypothesis we arrived at in our previous work showing DNA condensation in the presence of *Vc*DciA, suggesting that DciA could be a DNA chaperone by intercalating itself between the two DNA strands to stabilize it^13^. *Vc*DciA may thus constrain the mobility of the DNA.

### CapFlex (FIDA) analysis of DciA LLPS confirmed the importance of *Vc*DciA and DNA concentrations

For further comprehension of *Vc*DciA LLPS formation, we analyzed the droplets in different conditions with a FIDA (Flow-Induced Dispersion Analysis) device. CapFlex was recently used for the thermodynamic and kinetic characterization of protein LLPS ^26^. In these experiments, samples flowed through a capillary. When a droplet passes the detector it will register a spike signal (see Methods). Thus, the number of spikes is correlated to the total number of droplets, and the signal intensity of the spikes is proportional to the size of the droplets. It is noted that the population of the free and of the bounded state of probe His Lite™ OG488-Tris NTA-Ni inside and outside the droplets are not known, because the link to the His-Tag of *Vc*DciA is non-covalent. The increase in spike confirms that both *Vc*DciA and the probe His Lite™ OG488- Tris NTA-Ni enter the droplets (Fig. 3A). In this assay and due to the non-covalent labeling, the degree of phase separation is quantified solely by the number of spikes and their relative intensity and not the baseline intensity as demonstrated by Stender et al. ^26^ . Therefore, the partitioning of the label into the droplets should not impact the conclusion.

**Figure 3.**
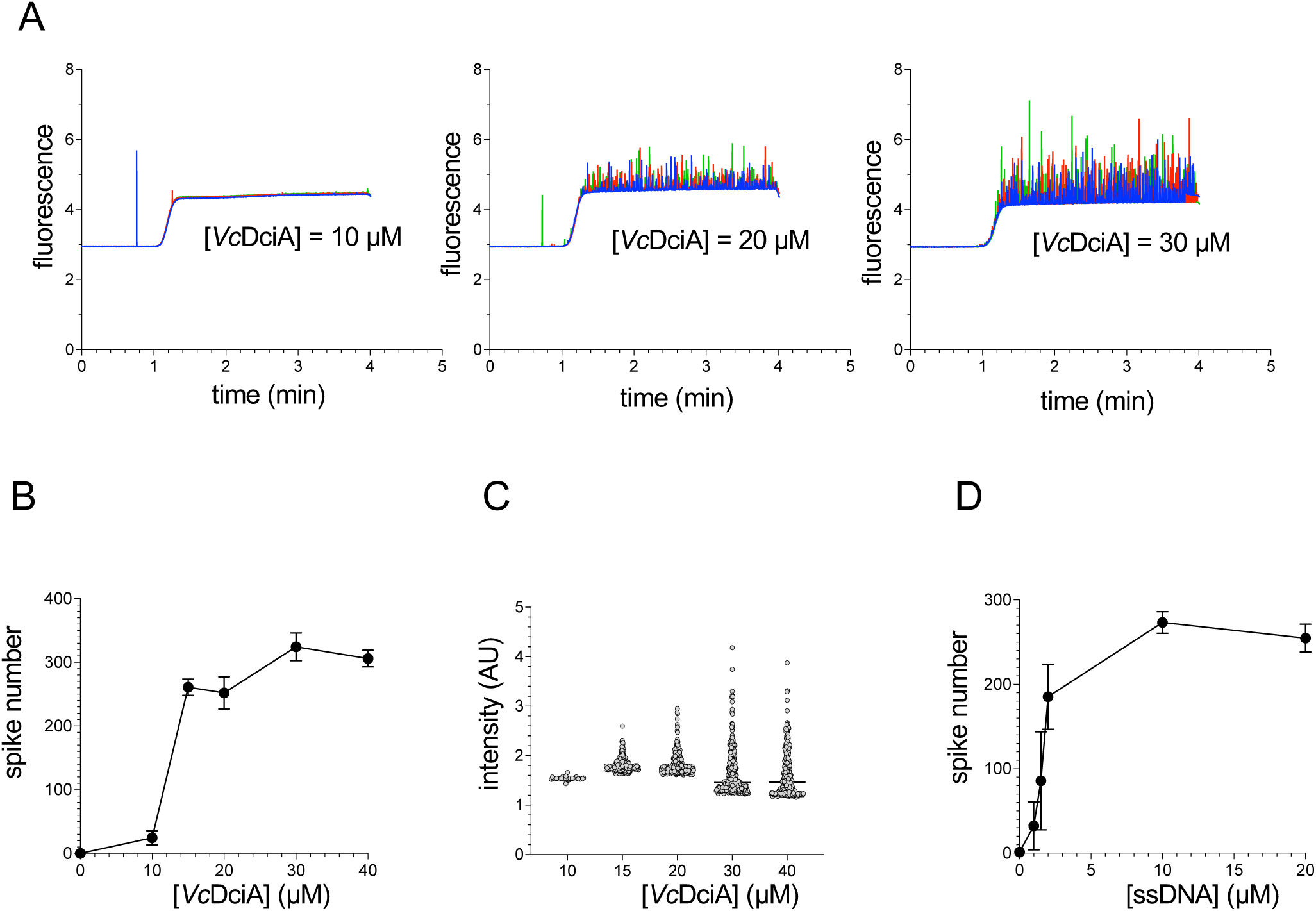
CapFlex (FIDA) analysis of LLPS. **A.** Signal spikes observation. Three experiments are presented in the constant presence of poly-dT50 (10 µM) with *Vc*DciA at the indicated concentration. For each concentration, three measurements (triplicates) are reported with three different colors (blue, red, and green). **B.** Spikes counting in various amounts of *Vc*DciA. Quantification of spikes number was performed on triplicate experiments and is reported for different *Vc*DciA concentrations, in the constant presence of poly-dT50 (10 µM). **C.** Spikes intensity report in various amounts of *Vc*DciA. Each droplet intensity is presented for different *Vc*DciA concentrations. **D.** Spikes counting in various amounts of ssDNA. Quantification of spikes number was performed on triplicate experiments and is reported for different poly-dT50 concentrations, in the constant presence of *Vc*DciA (20 µM).

Using this technology, we tried to see the droplets formation/generation as a function of time and protein concentration, in similar experimental conditions as we observed LLPS under the microscope. We first made experiments at a fixed concentration of ssDNA (10 µM), with increasing concentrations of *Vc*DciA (Fig. 3A and 3B). Droplets were analyzed in triplicates. We can see that the spikes, almost absent at 10 µM of *Vc*DciA, increase in intensity depending on the concentration of *Vc*DciA, i.e. the peaks are higher at 30 µM than at 20 µM (Fig. 3A). But from 15 µM their number remains very stable despite the increase of the protein concentration (Fig. 3B). This suggests that at a certain concentration of *Vc*DciA, no more new drops are formed, but their size might increase. This is confirmed by the observation of the spreading of the droplet intensity distribution (Fig. 3C) which reflects an increase in the average droplet size, even if we can’t exclude that it could be an increase in the concentration of labeled *Vc*DciA inside the droplets ^26^.

It is concordant with what we observed in the photonic microscopy experiments. Larger droplets can be observed as the concentration of *Vc*DciA increases. This phenomenon was even more obvious if we let them settle for 1 hour, probably due to the additional effect of droplet fusion (Fig. 2A and Supplementary Fig. 1D).

The impact of the DNA in the formation of the droplets has been studied by maintaining the *Vc*DciA concentration constant at 20 µM and by varying the concentration of the ssDNA (Fig. 3D). We confirmed that LLPS formation is strictly dependent on DNA because there are no spikes in the absence of DNA, but that little DNA is needed (2 µM) to reach a large number of spikes. The number of spikes varies only marginally afterward, when the DNA concentration is increased.

### EM and CryoEM analysis *of Vc*DciA ssDNA complexes : from networks to LLPS formation

Our previous EM analysis of *Vc*DciA/DNA interactions, even at low protein concentration, suggested properties of *Vc*DciA to condensate ssDNA ^13^. We have explored these properties with 50 nucleotides long poly-dT and with large ssDNA provided from virion PhiX174. The positive staining associated with the darkfield image mode is a powerful and relevant method to analyze such behavior, as previously shown for HIV nucleo-capsid protein ^27^ and confirmed as LLPS ^28^. These complexes were visualized in parallel with negative staining.

*Vc*DciA/poly-dT complexes form small spheroid complexes for 250 nM *Vc*DciA and 1.5 nM poly-dT. Their sizes increase from 20 to 50 nm for 250 nM *Vc*DciA (Fig. 4A a1-4) to 200 nM and more for 1 and 2 µM (Fig. 4A c-d). Some of these spheroids appear to grow by particle fusion mechanisms (Fig. 4A a2, b1, b3, c1, d1-3).

**Figure 4.**
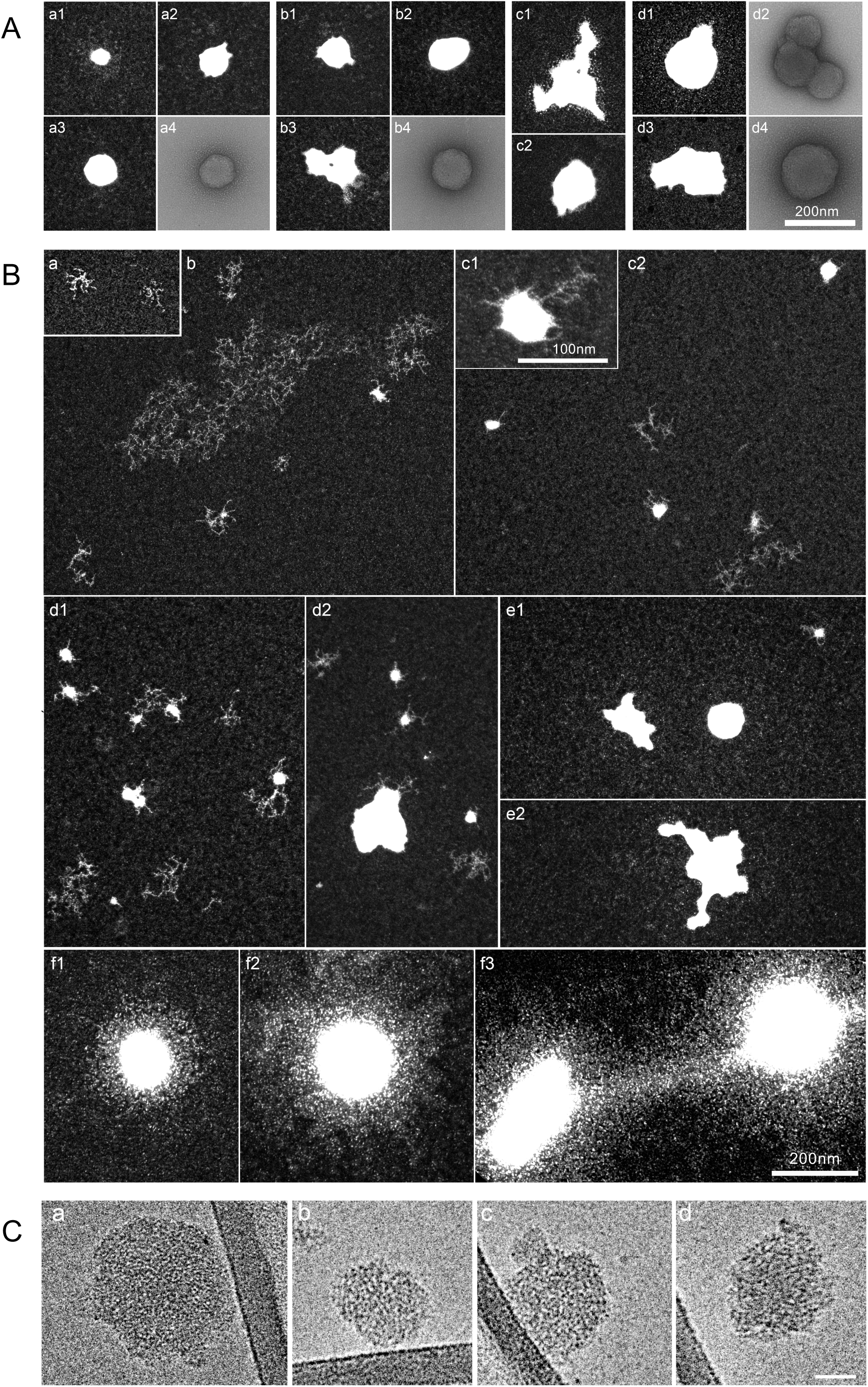
Electron microscopy and CryoEM analysis of *Vc*DciA-DNA complexes. **A.** LLPS formation in the presence of poly-dT50 oligonucleotide (1.5 µM), with 250 nM (a1-4), 500 nM (b1-4), 1 µM (c1-2) and 2 µM (d1-4) of *Vc*DciA. Scale bar : 200 nm. **B.** LLPS formation in the presence of PhiX174 virion ssDNA genome (1.5 µM) without (a) and with 250 nM (b, c1-2), 500 nM (d1-2), 1 µM (e1-2) and 2 µM (f1-3) of *Vc*DciA. Scale bars : 200 nm and c1 : 100 nm. All complexes were obtained by positive staining method visualized in darkfield mode except a4, b4, d4 where samples were obtained by negative staining visualized in brightfield mode. **C.** a-d CryoEM of LLPS resulting from poly-dT50 (10µM) – DciA (14µM) interactions. Scale Bar : 50 nm.

Interaction between PhiX174 ssDNA genome and *Vc*DciA leads to the formation of DNA networks at 250 nM *Vc*DciA as well as small spheroids (Fig. 4B b). These figures appear frequently under the same conditions (Fig. 4B c1-2), with structures of around 50 nm in diameter. Growth by fusion of these structures appears more clearly at 500 nM *Vc*DciA, producing spheroids of 50 to 200 nm (Fig. 4B d1-2). These growth mechanisms continue at 1 µM, where the structures appear irregularly shaped (Fig. 4B e1-2), whereas at 2 µM the shapes become more regular, with sizes of the order of 200 nm and more (Fig. 4B f1-3).

EM allows us to characterize the formation and growth stages of these LLPS, which occur from 250 nM *Vc*DciA upwards. We clearly show that such condensation results from network formation induced by bridging events, mediated by “DNA-protein”-“protein-DNA” interactions (Fig. 4B b, d1-2, f3). It should be noted that such observations become impossible at concentrations above 2 µM, as the carbons on which the samples are deposited do not resist, probably due to the large size and weight of the LLPS.

In order to study these networks at a more precise level of characterization, we have investigated the structure of *Vc*DciA-poly-dT complexes by cryoEM (Fig. 4C). We obtained similar spherical condensates of 50 to 200 nm with variations in sizes and shapes. The aim of such approaches was also to decipher the organization within the condensates and not only a 3D reconstruction of the spheroids. Our observations suggested a well-organization without being able to define it precisely. The heterogeneous LLPS condensates represent the various stages of the condensation process. DNA networks yield irregular condensates, followed by the formation of more compact spherical structures that grow into larger spherical aggregates by fusion with other spherical condensates.

Although our LLPS reconstituted for the confocal microscopy experiments are often larger than a bacteria, the spheroids analyzed with nanometric EM resolution have sizes compatible with the bacteria diameter usually close to 1 µm. The low concentration of 250 nM of DciA, sufficient to form LLPS, suggests that these structures may exist *in vivo*, probably to create a non-membrane compartment in the proximity of the *oriC* or arrested replication forks, to facilitate replication initiation and restart.

### *Vc*DciA recruits *Vc*DnaB into the droplets

The formation of LLPS by *Vc*DciA in the presence of ssDNA is similar to previous studies reported for replication initiation factors in eukaryotes^20^. Indeed, it was demonstrated that the proteins involved in eukaryote replicative helicase loading, ORC, Cdc6, and Cdt1 contain intrinsically disordered regions (IDRs) that drive LLPS ^29^. Although these proteins have no structural similarity with *Vc*DciA, their function is similar. It was also demonstrated that the replicative MCM helicase can be recruited in those LLPS ^29^. We then decided to investigate for an analog function for *Vc*DciA.

We previously demonstrated by SPR and BLI that the loading on the ssDNA of the replicative helicase, from *Vibrio cholerae* (*Vc*) as well as from *E. coli* (*Ec*), is increased by *Vc*DciA^11,15^. We then tried to see if *Vc*DciA could attract/recruit *Vc*DnaB into preformed LLPS (Fig. 5A). We performed this experiment in the presence of ATP and MgCl2, which are required for helicase hexamerization and activity.

**Figure 5.**
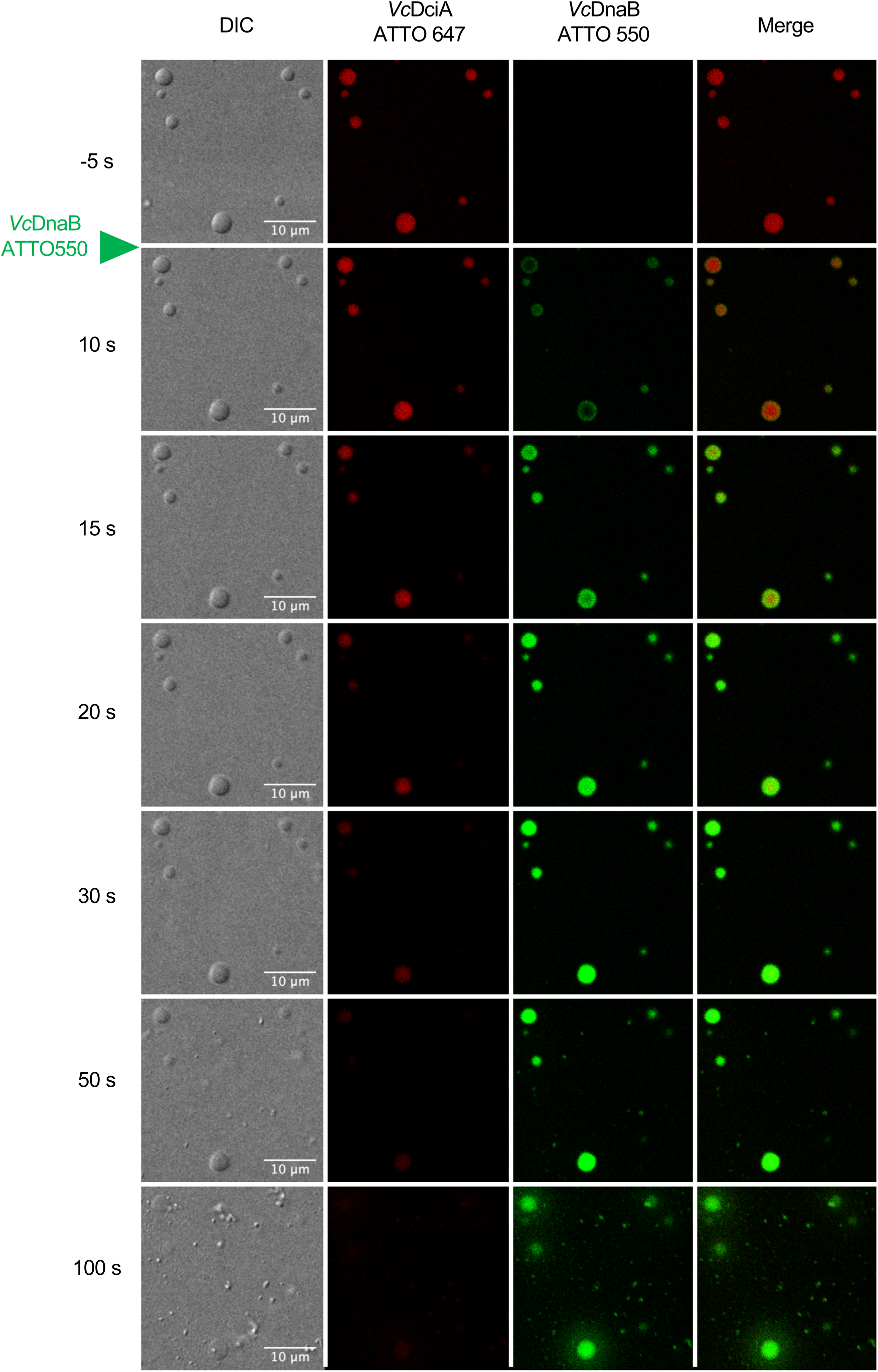
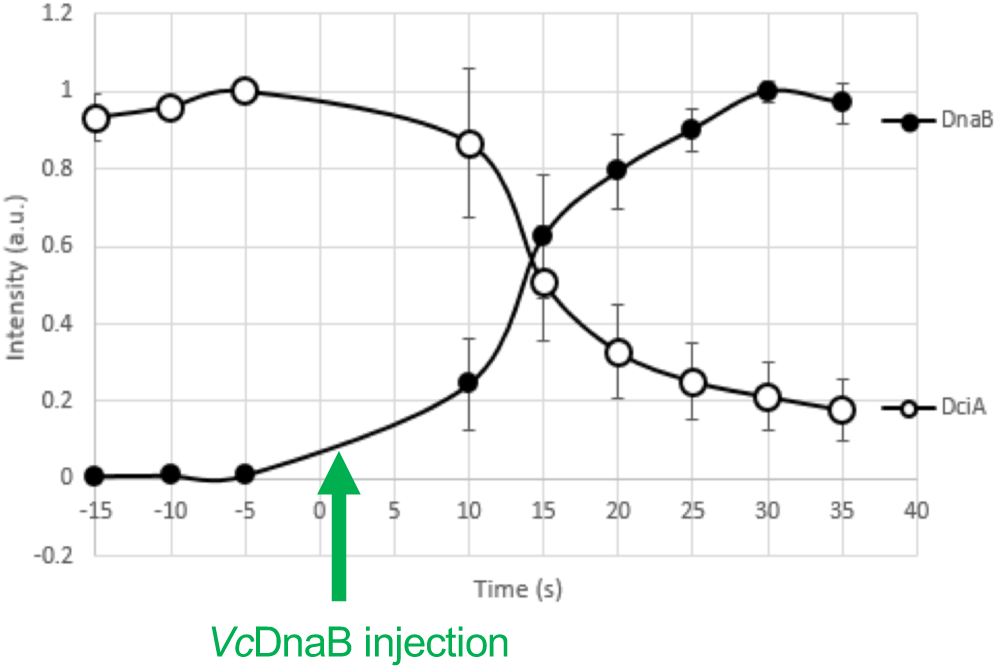
*Vc*DciA recruits *Vc*DnaB into the droplets. LLPS were formed with 12 µM of *Vc*DciA-ATTO 647 in the presence of 10 µM of poly-dT50 in MN buffer containing 1 mM ATP with 10 mM MgCl_2_. Between 0 and 10 seconds, *Vc*DnaB-ATTO 550 was added at the final concentration of 4 µM. A. Representative confocal images. Images collected at indicated times were presented as they illustrated the different events. 10 µm scale is indicated. B. Quantification. Representative graphic of the average of the mean intensity (± standard deviation) within a droplet, normalized to 1. Full circles represent *Vc*DnaB-ATTO 550 and open circles *Vc*DciA-ATTO 647. The average was measured on n=6 distinct droplets.

We first formed droplets with *Vc*DciA (labeled with ATTO 647) and ssDNA (unlabeled), before adding (at t = 0 second in Fig. 5A) *Vc*DnaB (labeled with ATTO 550). We observed that *Vc*DnaB rapidly locates on the periphery of the droplets, and is quickly fully internalized. In Fig. 5B we quantify the normalized droplet intensity for both labels. This approach confirmed that the fluorescence intensity of *Vc*DnaB starts to increase progressively inside the droplets 15 seconds after its addition, and reaches a plateau after 40 seconds. In parallel, the *Vc*DciA signal inside the droplets decreases with an inverse behavior, until it stabilizes at around 20% of its initial concentration. At 100 seconds, droplets are destructured as can be seen in the DIC image, and small dense satellites appear, while the fluorescence of *Vc*DnaB persists, with a blurred halo around the location of the restructured droplets (Fig. 5A). To ensure that the fluorescent signal observed in the droplets is due to DnaB-ATTO 550 and not to the binding of the free dye to *Vc*DciA, we verified that when Ni-NTA-ATTO 550 is added alone on *Vc*DciA/DNA droplets, Ni-NTA-ATTO 550 is never recruited in the LLPS (Supplementary Fig. 3A).

In the presence of *Vc*DnaB, the decrease of *Vc*DciA-ATTO 647 correlates with the fast release of *Vc*DciA during the helicase loading experience described previously in BLI assays ^11^, suggesting that *Vc*DciA is well ejected from the droplets. In the experimental conditions developed in this study, *Vc*DnaB alone is not visible since it remains soluble (Supplementary Fig. 3B). If we mix together *Vc*DnaB with *Vc*DciA and ssDNA, without first preforming the *Vc*DciA/ssDNA LLPS, we observed undefined structures that could correspond to precipitates with both fluorescent tags inside (Supplementary Fig. 3C). This aggregation indicates that the proteins precipitate together in the presence of ssDNA in these conditions, as we have often observed when preparing the ternary complex samples for structural analysis by SAXS or crystallization. To this date, the underlying cause of this precipitation remains unknown. However, it is plausible that the simultaneous mixing of the three partners may hinder proper complex organization. The sequential addition of partners, i.e. protein-protein-DNA or protein-DNA-protein, is then probably important, and may reflect the sequentially of DnaB recruitment by DciA at *oriC*.

### *Ec*DnaC from *Escherichia coli* shows a behavior close to that of *Vc*DciA

There is no sequence or structural similarity between DciA and DnaC/I. While *Vc*DciA consists of a KH-like domain that binds nucleic acids, followed by a non-folded C-terminal extension that structures into 2 small α-helices upon contact with DnaB ^15^, *Ec*DnaC consists of an AAA+ domain preceded by a long α-helix that contacts DnaB ^4^. Nevertheless, both loader proteins target the same surface on DnaB, the LH/DH module, which may allow us to qualify this similarity as an evolutionary convergence ^17^.

Here, we observed that *Ec*DnaC is also able to form droplets with ssDNA, in the protein concentration conditions set up for *Vc*DciA (Fig. 6A). We can observe that there is indeed phase separation, but that the droplets generally appear smaller than those formed by *Vc*DciA (Supplementary Fig. 4A). Furthermore, we do observe contacts between droplets that tend to end up in incomplete fusion events, or into droplets that do not relax into single spherical (homogenous) droplets (Supplementary Fig. 4B). This suggests that LLPS formed by *Ec*DnaC are less fluid than those of *Vc*DciA are. The fact that they do not fuse shows that they are viscous, and more of a solid than a liquid phase.

**Figure 6.**
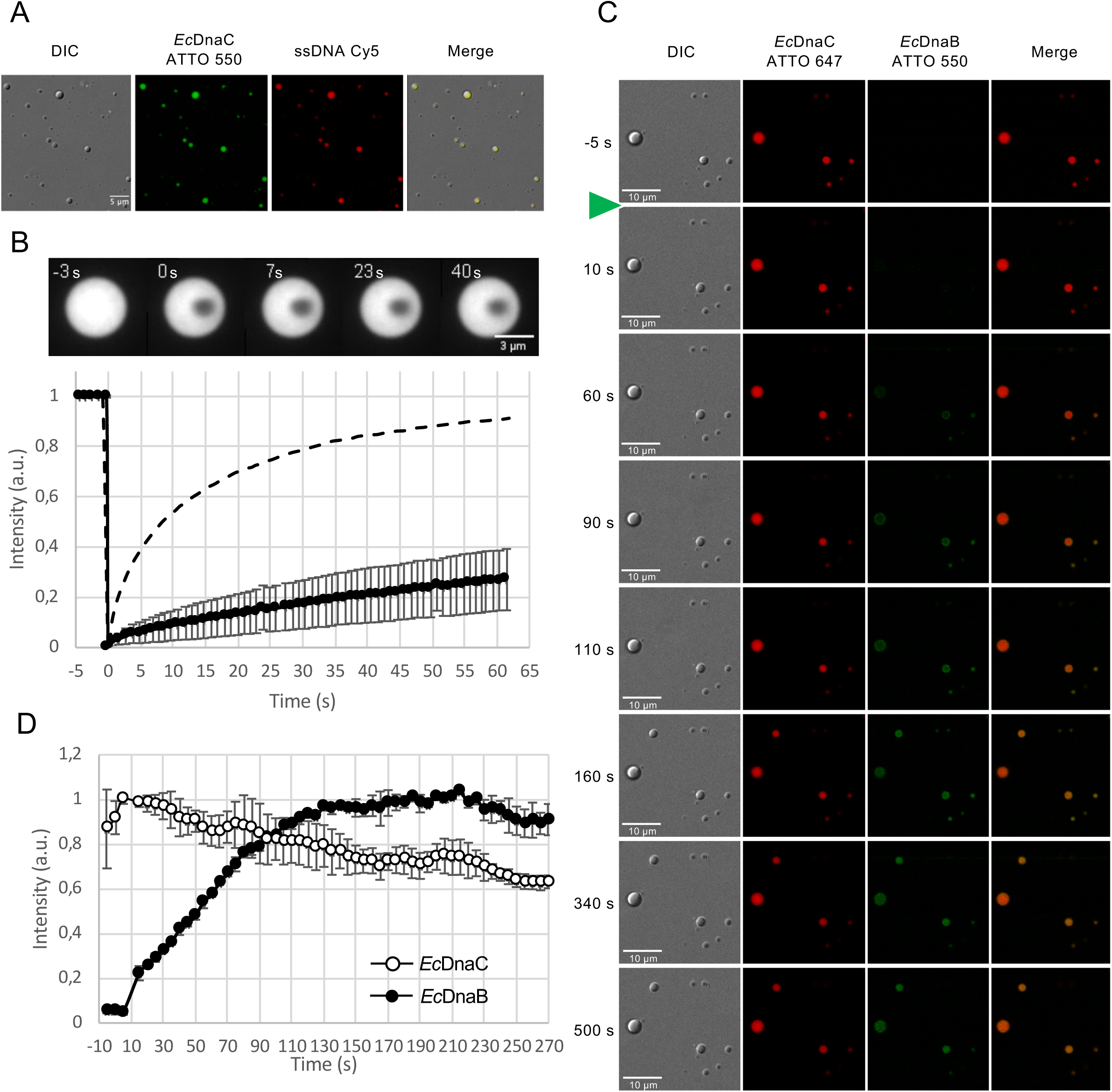
Microscopy analysis of droplets formed by *Ec*DnaC. **A.** Representative confocal image of *Ec*DnaC labeled in the presence of fluorescent ssDNA. LLPS were formed with 15 µM of *Ec*DnaC labeled with ATTO 550, in the presence of 10 µM of poly-dT50 and 0.2 µM of ssDNA labeled with Cy5 (ssDNA-Cy5). 10 µm scale is indicated. **B.** FRAP experiments on *Ec*DnaC droplets. LLPS were first formed with 30 µM of DnaC-ATTO 550 in the presence of 10 µM of poly-dT50. On top, representative time-lapse imaging of a FRAP experiment. FRAP was applied on a central sub-region (1 μm diameter). Time is indicated in seconds. The bleaching event is performed at time 0s. The scale bar is 3 μm. On the bottom part is a representation of the mean fluorescence intensity recovery of FRAP experiments presented on the top (±Standard deviation). Data were obtained from 3 independent replicates (n=25). The dotted line is a reminder of the mean *Vc*DciA recovery (shown in comparison). **C.** *Ec*DnaC recruits *Ec*DnaB into the droplets. LLPS were formed with 12 µM of DnaC-ATTO 647 in the presence of 10 µM of poly- dT50 in MN buffer containing 1 mM ATP with 10 mM MgCl_2_. 5 µM scale is indicated. Between 0 and 10 seconds, *Ec*DnaB-ATTO 550 was added at the final concentration of 4 µM. Images collected at indicated times were presented as they illustrated the different events. **D.** Quantification. Representative graphic of the average of the mean intensity (± standard deviation) within a droplet, normalized to 1. Full circles represent *Ec*DnaB-ATTO 550 and open circles *Ec*DnaC-ATTO 647.

Nevertheless, we can observe that the ssDNA co-localizes with *Ec*DnaC (Fig. 6A). To better understand the nature of those droplets we performed FRAP experiment in the same condition as for *Vc*DciA (Fig. 6B). Analysis of the FRAP experiments showed that only 26.68±12% of *Ec*DnaC associated fluorescence recovered after 1 minute, with a 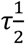 of 47.2±16 s. This is much slower than the recovery observed for *Vc*DciA. Moreover, the FRAP experiments on *Ec*DnaC showed a linear recovery of the fluorescence, which does not reflect LLPS behavior. This result and the observation that the droplets rarely fuse and do not relax after fusion, lead us to the conclusion that *Ec*DnaC droplet do have not a liquid-like nature but rather forms a gel-like object. To gain more insight into the interactions inside *Ec*DnaC droplets, we performed again FRAP experiments on *Ec*DnaC-ATTO 550 and ssDNA-Cy5 at the same time on the same droplet with the same intensity (Supplementary Fig. 4C-D). The ssDNA is still slower than the protein inside droplets as with *Vc*DciA. Nevertheless, the ssDNA has a mobile fraction of 24.6±6.7% with a 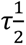 of 84.3±21 s which is even slower than in *Vc*DciA-ssDNA droplets. This is consistent with what we could expect inside gel-like droplets. We can observe the fact that the reduction of the recovery of the protein included in the droplet reduces the recovery of the same ssDNA too. This confirms the Protein-DNA interactions inside the droplets.

We were also able to see the recruitment of *Ec*DnaB into the droplets, but with a much slower speed than in the case of *Vc*DciA (Fig. 6C). While *Vc*DnaB is visible at the periphery of LLPS as early as 10 s, it takes more than 1 min for *Ec*DnaB to be visible at the periphery of *Ec*DnaC LLPS. It takes at least 340 s for *Ec*DnaB to be incorporated into the droplets (*vs* 20 s for *Vc*DnaB into the *Vc*DciA droplets), and no disappearance of *Ec*DnaC was observed up to 500 seconds whereas *Vc*DciA was ejected in 50 seconds (compare Fig. 5 to Fig. 6C). Indeed, we observe a slight decrease of the DnaC-ATTO 550 fluorescence intensity inside the droplets (Fig. 6D). This difference of behavior is probably due to the more stable, more fixed state of the gel-like *Ec*DnaC/ssDNA droplets.

### Concluding remarks

DciA is the replicative helicase loader predominantly found in the bacterial world. Contrary to what is known for DnaC, the loader that replaced DciA late in evolution ^10^, the mechanism that DciA applies to the helicase to ensure its loading remains to be understood, as the structure of the DnaB•DciA complex that we have solved by crystallography shows no impact on the conformation of DnaB by DciA ^11,30^. It is therefore likely that a further player is required to ensure this fundamental function. We postulate that this other partner could be DNA itself.

DciA, with a very electropositive overall charge, has the structural characteristics required for a DNA condenser, since it is composed of a globular KH-type domain described as interacting with RNA or DNA, and an unfolded extension. Hence the interest in exploring the relationship between DciA and DNA, which have together the property of forming liquid-liquid phase separations. This paper is dedicated to the characterization of these LLPS, using photonic and electron microscopy approaches. We have characterized their appearance, their size and fluidity, their ability to fuse, and their capacity to recruit DnaB. Positive staining electron microscopy images and cryoEM analysis are providing evidence that the condensates we observed result from network formation induced by bridging events. These bridges are formed by interactions mediated by “DNA-protein”-“protein-DNA” interactions.

These new data are consistent with the model we had arrived at, which proposed that DciA could be a chaperone for the DNA ^13^. Here, we show that DnaC also has the capacity to interact with DNA within LLPS, although these are gelled and much less fluid than those formed by DciA. Our FRAP experiments show that both proteins and DNA are slowed down when engaged in interactions in these droplets, proving their contacts, which we have shown to be partly due to electrostatic interactions.

We can therefore propose a certain universality in the mechanism of helicase recruitment by its loaders, even if each one seems to have its own specificities. It was previously demonstrated that eukaryotic replication initiators can recruit the replicative MCM helicase in LLPS, and that condensation in the nucleus could be an important regulatory mechanism for DNA replication ^29^. The existence of non-membrane compartments in bacteria seems to be possible with helicase loaders as drivers, and may therefore be a conserved phenomenon. Our results and the discovery of this ability of DciA - to a lesser extent DnaC - to form LLPS enable us to propose that the recruitment of DnaB into the cell takes place sequentially, starting with the formation of a non-membrane compartment by DciA in the environment of genome zones such as *oriC* or arrested replication forks. These areas would facilitate the recruitment of DnaB to the ssDNA on which it is to be loaded, resulting in the release of DciA and the initiation of replication or the restart of replication following replisome reassembly. The search for these non-membranous compartments in bacteria is therefore becoming a priority, in order to demonstrate the reality of their existence.

## Methods

### Oligonucleotides used

The poly-dT50 oligonucleotide (repetition of 50 thymines) was mixed with proteins for LLPS formation observed by photonic microscopy. The 50 nucleotides long oligonucleotide Oso26 (5’-CCAGGAATACGGCAAGTTGGAGGCCGGGCTGGATGGAGACTAAGCTTTGG-3’) is labeled at its 5’ extremity by a cyanine-5 (Cy5) and was mixed to follow DNA during fluorescent LLPS imaging. A 50 nucleotides long oligo-dT is used for the electron microscopy experiments, as well as the ssDNA of the PhiX174 genome (New England Biolabs).

### Protein samples preparation

*Vc*DnaB, *Ec*DnaC and *Ec*DnaB were all 6His-tagged at the C-terminus during the cloning process while *Vc*DciA is tagged at the N-terminus. After over-expressed in the *E. coli* Rosetta(DE3)pLysS or the BL21-Gold(DE3) strains, they were purified as described previously^11,30^. Briefly, lysis were performed in buffer A (NaCl 200 mM, Tris–HCl 20 mM (pH 7.5)) by sonication and the His-tagged proteins were purified on a Ni-NTA column (Qiagen Inc.), eluted with imidazole in buffer A but complemented by ATP 1 mM + MgCl2 3 mM in the case of the *Vibrio cholerae* helicase. *Vc*DciA or *Ec*DnaC were then loaded onto a Heparin column in a phosphate buffer (NH2PO4 20 mM, NaCl 100 mM (pH 5.8)) and eluted with a gradient of NaCl (from 0.1 to 1.5 M). The two helicases were loaded onto a Superdex TM200 column 16/600 (GE), equilibrated against NaCl 100 mM, Tris–HCl 20 mM (pH 8.8) complemented by ATP 1 mM, MgCl2 3 mM in the case of *Vc*DnaB. Their purification was completed with a MonoQ column (Amersham Pharmacia Biotech) in Tris–HCl (20 mM, pH 8.8) and a gradient of NaCl (from 0.1 to 1 M).

### Turbidity analysis of LLPS formation

LLPS was monitored by turbidity (optical density at 620 nm) by an Infinite-200-PRO- Fluorimeter (Tecan, Mannedorf, Switzerland). Analysis was performed at 30°C in 50 µl of the indicated buffer. Protein concentrations are indicated in the figures.

### Protein Labelling for microscopy imaging

His-tagged proteins were labeled using Ni-NTA-ATTO 550 or Ni-NTA-ATTO 647 from Merck Group. Proteins at a final concentration of 300 µM were incubated 15 minutes on ice in 20 µl of MN buffer (50 mM MES pH6, 150 mM NaCl) with the indicated dye at a concentration of 30 µM (ratio dye/protein of 1/10).

### Glass treatment for LLPS Microscopy Imaging

Two kinds of coating were tested for LLPS visualization: pluronic or neutral coating (siliconizing). For Pluronic coating, glass surfaces were treated by adding a filtered 10% solution of Pluronics F-127 and incubated for ∼1h. The chambers were rinsed 4-5 times with MilliQ water and remain immersed in water until use. For neutral coating, glass surfaces were treated first with siliconizing solution Sigmacote (from Sigma) for 20 seconds, washed 5 times with MilliQ water, and treated after with Pluronics as in case of Pluronic coating.

### DIC and Fluorescent Microscopy Imaging

Droplets of protein were formed in MN buffer with 10 µM of a poly-dT50 nt single-stranded oligonucleotide. Samples (20 µl) were loaded on a µ-Slide 18 Well Glass Bottom (Ibidi, GMBH). Images were acquired on a LEICA SP8X inverted confocal laser scanning microscope equipped with a 63x HC Plan Apochromat CS2 oil-immersion objective (NA: 1.4) (Leica) with hybrid GaAsP detectors (Hamamatsu). For DIC images, we used the 481 nm wavelength. A white light laser was used to excite ATTO 550 at 550 nm and both ATTO 647 and Cy5 at 648 nm. The fluorescence is collected between 560 and 601 nm for ATTO 550, and between 675 and 726 nm for both ATTO 647 and Cy5. Image treatment and analysis were performed using FIJI free software ^31^.

### FRAP experiments

FRAP acquisitions were performed on an inverted Eclipse Ti-E (Nikon) microscope coupled with a CSU-X1-A1 Spinning Disk (Yokogawa), a 100x Apochromat TIRF oil immersion objective (NA: 1.49) (Nikon) and a Prime 95B sCMOS camera (Photometrics). The whole system is driven by MetaMorph software version 7.7 (Molecular Devices), and iLas 2 module (GATACA Systems) for FRAP.

All the bleaching events are performed in a circular region with a diameter of 1 μm and monitored over time for fluorescence recovery using a Quad band 440/40 nm, 521/20 nm, 607/34 nm, 700/45 nm emission filter (Semrock). For the *Vc*DciA-ATTO 550 FRAP experiment, we used the 561 nm, 100 mW laser (Coherent) set at 4% for the imaging and 44% for the bleaching, with an exposure time of 200 ms. The FRAP sequence was composed of a short pre-bleach sequence of 3-5 s (1 image/s), the bleaching event, 2 successive post-bleach sequences of 20 s (2 images/s), and then a last sequence of variable length (1 image/s).

For the dual colors FRAP we adjusted the laser power to obtain similar bleaching amplitude in both channels (but this adjustment induces a variability between the recovery of the experiments done with one color than those done with 2 colors as the intensity isn’t the same). To get rid of most of the manipulation errors, we FRAP the DNA marked and *Vc*DciA marked at the same time on the same droplets and then calculated the ratio between each half-time recovery 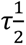. To avoid acquisition-related bleaching, ssDNA is acquired at a frequency 5 times lower than that used for the protein, and the exposure time was 200 ms for *Vc*DciA and 100 ms for the ssDNA. The FRAP sequence was composed of : 5 s pre-bleach (1 image/s), bleaching event (150 ms), 2 successive post-bleach sequences of 20 s (1 image/s) and then 180 s (1 image/3 s).

FRAP curves were independently corrected and processed to obtain a double normalization as follow : the mean intensity of every bleached region was measured and the background intensity was subtracted by measuring a region outside the droplet. Acquisition-related bleaching correction is performed by dividing values by the whole droplet fluorescence intensity (as the bleached region can not be considered as negligible). Then to display the recovery curves from 0 to 1, a double normalization is performed using the average of the pre-bleached signal and the 1st post-bleached value.

The mean indicated in the figures is the one calculated from all the measurements pooled. Custom-built macro was written in ImageJ to perform quantitative image analysis. As the recovery curves display a biphasic aspect, the mean curve was fitted by a double exponential function to extract both half-time recovery 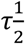 and the mobile fraction.

When we were able to image and record the recovery to the plateau, the value of the plateau after fitting was used for the calculations. Otherwise, when the plateau was not achieved, we used the last recorded value to avoid fitting artifacts at the infinity.

### FIDA experiments

For FIDA analysis, His tagged proteins were labeled using Ni-NTA-ATTO 488 (Merck Group) at a ratio of ATTO/protein = 1/500 in MN buffer (50 mM MES pH 6, 150 mM NaCl).

Capflex was performed with a non-covalent Probe His Lite™ OG488-Tris NTA-Ni Complex unlike the covalent labels used before ^26^. This means that the baseline corresponds to the concentration of Probe His Lite™ OG488-Tris NTA-Ni Complex in the dilute phase and not DciA. However, the label still enters the droplets and the signal can thus be used as a means of quantification. The Capflex method was described in detail previously ^26^. Briefly, a sample is flowed through a capillary of 1 mm with an inner diameter of 75 µm. The auto- sampler and the capillary chamber is temperature controlled to a precision of 0.1 °C. As the sample passes the confocal fluorescence detector the baseline corresponds to the dilute phase concentration of the fluorescent probe and if the fluorescent probe enters the droplets, each time a droplet passes the detection window a spike is observed. This means the total number of spikes correlates to the total number of droplets ^26^.

For the measurements of droplet formation, protein samples were prepared in MNP buffer (50 mM MES pH 6, 150 mM NaCl, 0.1% Pluronics F-127) containing or not poly-dT50 nt single-stranded oligonucleotide. Proteins at the indicated concentration were incubated in the sample trays at 25 °C, and the capillary chamber was kept at 25 °C. A standard capillary with an inner diameter of 75 μm and a length of 1 mm was used. The following set of instrument parameters was used : wash with NaOH 1 M for 120 s at 3500 mbar; equilibration with MNP buffer for 120 s at 3500 mbar; sample analysis in MNP buffer with or without DNA during 240 s at 650 mbar. Between 3 and 11 replicates were launched, with 9 minutes intervals between each sample.

Automated baseline and peak height detection was done using a script running in Jupyter 6.1.4 (available here: DOI 10.11583/DTU.14223116). For a detailed description please see ^26^.

### Electron microscopy

TEM observations were carried out with a Zeiss 912AB transmission electron microscope in filtered zero-loss darkfield or brightfield imaging mode. Electron micrographs were obtained using a Tengra digital camera (Olympus) and a Soft Imaging Software system (iTEM).

The positive staining associated with darkfield filtered imaging mode is usually used to analyze nucleoprotein complexes containing large DNA molecules when negative staining and brightfield filtered imaging mode are dedicated to protein structure. For positive staining, 5 µl of different mixed are deposited on hexagonal 600 mesh copper grids, previously covered with a thin carbon film and functionalized in a homemade device by glow-discharge in the presence of amylamine, providing NH^3+^ charge deposition onto the carbon surface. For negative staining, 5 μl of different mixed were adsorbed onto a 300 mesh copper grid coated with a collodion film covered by a thin carbon film, activated by glow-discharge to make the carbon film hydrophilic. In both cases, after 1 min, grids were washed with aqueous 2% w/vol uranyl acetate (Merck, France) and then dried with ashless filter paper (VWR, France). The nucleo-complexes were reconstituted without incubation at room temperature by mixing 250 nM, 500 nM, 1 μM or 2 μM of *Vc*DciA with PhiX 174 virion ssDNA or poly-dT oligonucleotides at a concentration of 1.5 μM nucleotides, in 50 mM MES buffer (pH 6) and 150 mM NaCl.

### Cryo-Electron microscopy

*Vc*DciA (14µM) /poly-dT (10 µM) LLPS were assembled at room temperature for 5 min in buffer (50 mM MES buffer (pH 6) and 150 mM NaCl). After a dilution of 10 in buffer, 3 µl of the mix were deposited on a glow-discharged Lacey grids (Agar scientific) and cryo-fixed using a EMGP2 (Leica) which has the particularity to one-sided blot sample (humidity 100%, temperature 18°C, blot time 1 s). Grids were imaged on a Tecnai F20 microscope (Thermo Fisher, USA) in low-dose conditions and operating at 200 keV and using a Falcon II direct detector (Thermo Fisher, USA).

## Acknowledgements

This work was supported by the French Infrastructure for Integrated Structural Biology (FRISBI) ANR-10-INBS-05, by funds from the Centre National de la Recherche Scientifique (CNRS), and by the Infrastructures en Biologie Santé et Agronomie (IBiSA). It has benefited from two I2BC facilities: Imagerie-Gif core facility, supported by the Agence Nationale de la Recherche (ANR-11-EQPX-0029/Morphoscope, ANR-10-INBS-04/FranceBioImaging; ANR-11- IDEX-0003-02/ Saclay Plant Sciences), and PIM facility supported by French Infrastructure for Integrated Structural Biology (FRISBI) (ANR-10-INBS-05). For cryoEM experiments, M. Nilges and the Equipex CACSICE (n° ANR-11-EQPX-0008) provided funding for the Falcon II direct detector. C.C. was supported by a Ph.D. fellowship from the French Ministry of Education.

## Data availability

The authors declare that the data supporting the findings of this study are available within the article and its Supplementary Information files, or are available from the corresponding authors upon request.

## Author contributions

S.M. and S.Q.-C. conceived this study; C.C., and S.Q.-C. purified the proteins; S.M., M.A-N. and M.N. performed the turbidity analysis; S.M., S.J. and R.L.B. performed the fluorescent microscopy imaging experiments; S.J. and R.L.B. performed the FRAP experiments; S.M. and E.GP.S. performed the FIDA experiments; E.L.C. and S.B. performed the electron microscopy experiments; S.B., H.W. and G.P-A. performed the cryo-electron microscopy experiments.

S.M. and S.Q.-C. wrote the paper with input from all authors.

