## supplementary figures for "The Bacterial Replicative Helicase Loader DciA is a DNA Condenser"

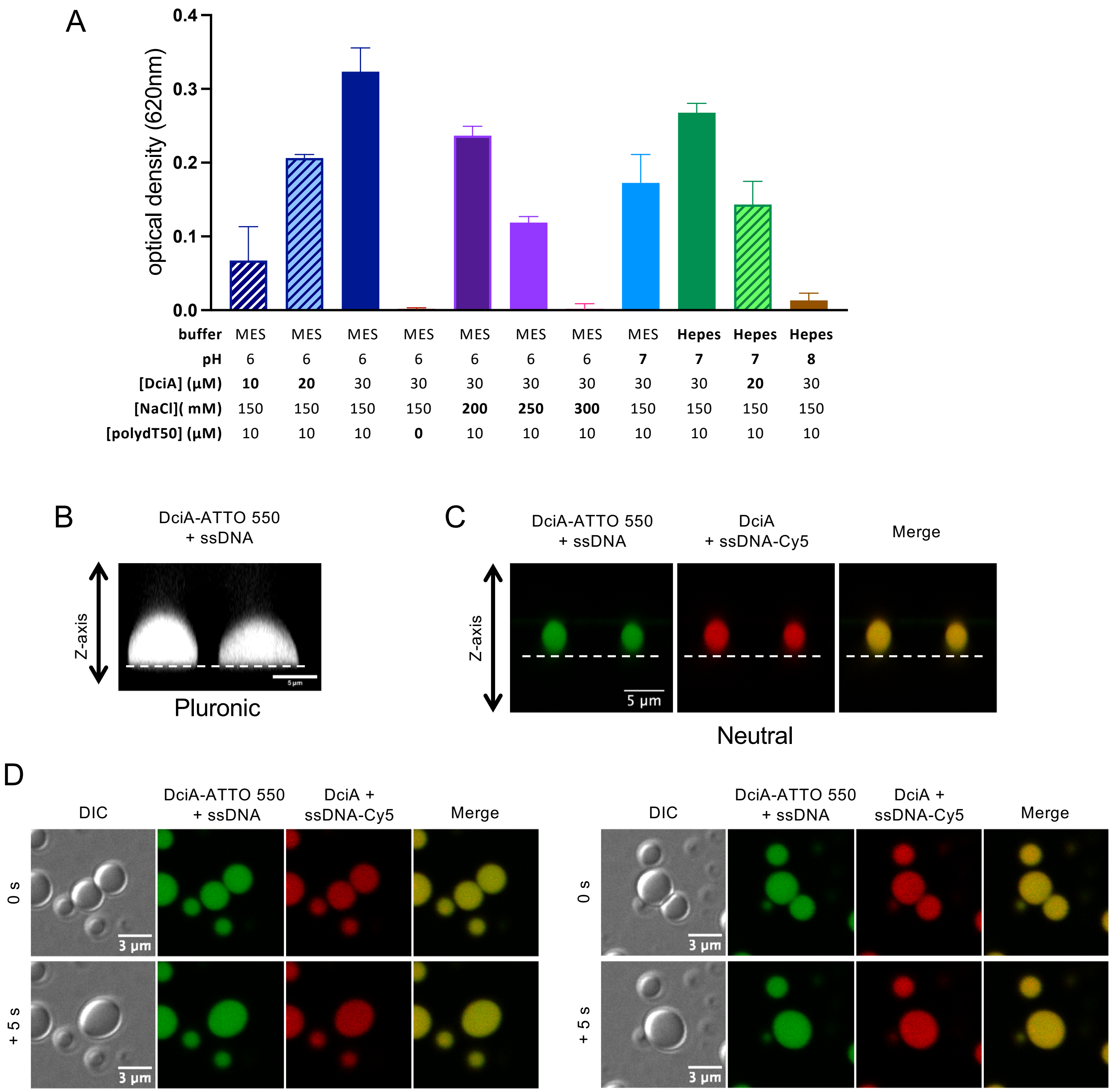


**Supplementary Figure 1: *Vc*DciA droplets analysis.**

**A.** Turbidity analysis of *Vc*DciA in solution in the presence of ssDNA. LLPS was monitored by turbidity (620 nm), in different buffers: MN pH6 (50 mM MES pH 6); MN pH7 (50 mM MES pH 7); HN pH7 (50 mM Hepes pH 7); HN pH8 (50 mM Hepes pH 8); in the presence of *Vc*DciA, NaCl, and poly-dT50 at the concentrations indicated in the table below the graph. Mean and standard deviation of three independent measures are reported.

**B.** 3D analysis of *Vc*DciA droplets on pluronic treated glass. Z wide image of two droplets was acquired with DciA-ATTO 550 droplets in the presence of poly-dT50. The dotted line corresponds to the position of the glass slide.

**C.** 3D analysis of *Vc*DciA droplets on neutral treated glass. Z wide image of two droplets was acquired with DciA-ATTO 550 droplets containing fluorescent ssDNA-Cy5. The dotted line corresponds to the position of the glass slide.

**D.** Example of DciA droplets fusion. Fusion after 40 minutes spreading of DciA-ATTO 550 droplets containing fluorescent oligonucleotide Oso26.

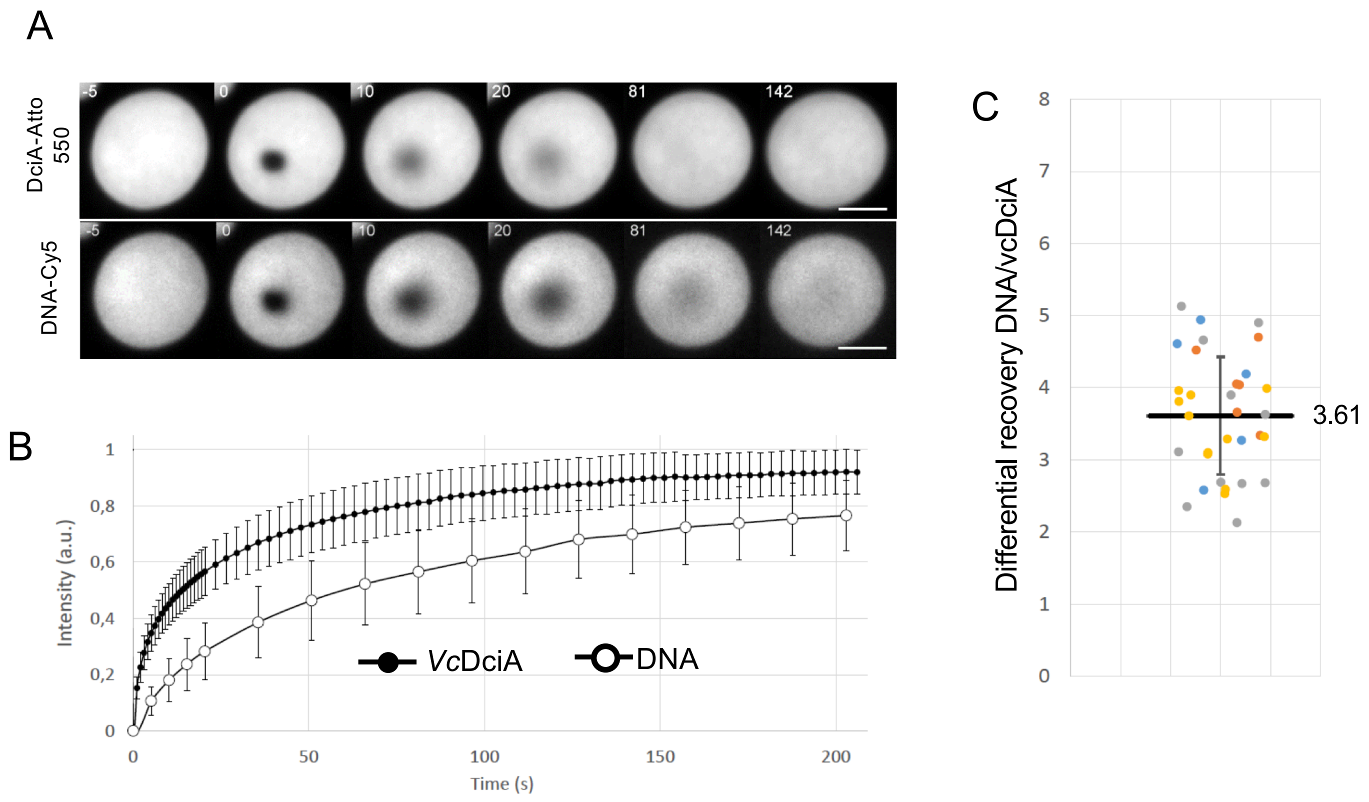


**Supplementary Figure 2 : DciA exhibits a higher mobility than DNA within a droplet.**

LLPS were first formed with 30 µM of DciA-ATTO 550 in the presence of 10 µM of poly-dT50 and 0.2 µM of ssDNA-Cy5. FRAP was applied on a central sub-region (1 µm diameter).

**A.** Time-lapse imaging of the same droplet. Top raw is the tagged protein and bottom raw is the tagged ssDNA. Time is indicated in seconds. The bleaching event is performed at the same time for both channels: time 0s. The scale bar is 3 µm.

**B.** Quantification of relative intensity recovered. Intensity recoveries of the mean fluorescence (± Standard deviation) of *Vc*DciA-ATTO 550 (full circles) and DNA-Cy5 (open circles) are presented. n=34 on four independent experiments.

**C.** Quantification of the difference in recovery between DNA and DciA. The ratio *τ1/2(DNA)τ12(DciA)* for each FRAP experiment to quantify the differential behavior at the droplet level and get rid of experimental variability is presented. The black horizontal line is the mean ratio (3.61± Standard Deviation), each color represents one independent experiment.

Supplementary Fig. 3 (legend on next page)

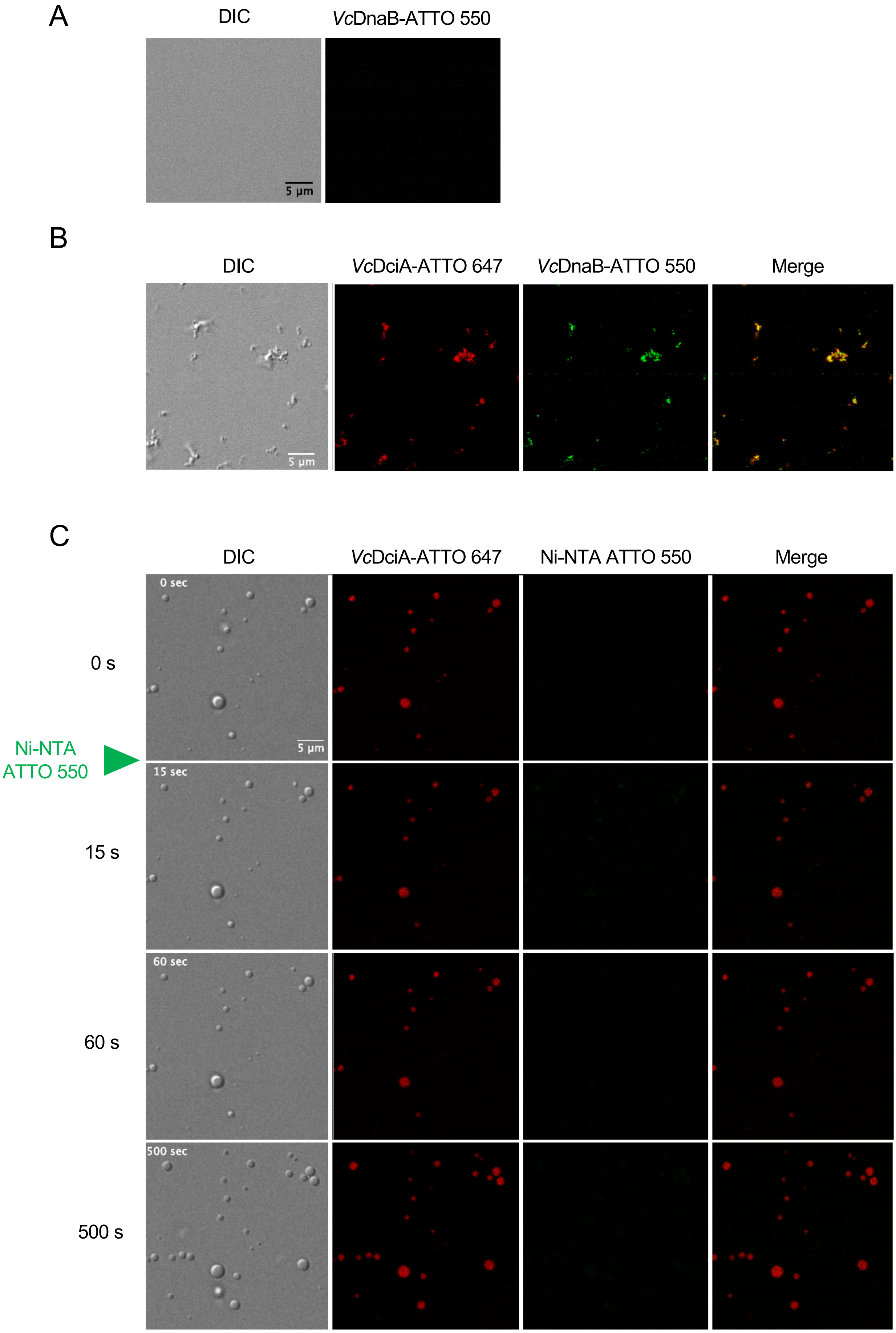

**Supplementary Figure 3 : Microscopy controls for *Vc*DnaB recruitment in *Vc*DciA-formed LLPS**

**A.** *Vc*DciA droplets do not recruit the dye in solution. LLPS were formed with 12 µM of *Vc*DciA-ATTO 647 in the presence of 10 µM of poly-dT50 in MN buffer containing 1 mM ATP with 10 mM $MgCl_2$. Between 0 and 15 seconds, Ni-NTA-ATTO 550 was added at the final concentration of 0.4 µM which correspond to the amount of Dye present in 4 µM of *Vc*DnaB-ATTO 550. Images collected at indicated times were presented.

**B.** Microscopy analysis of *Vc*DnaB-ATTO 550. *Vc*DnaB-ATTO 550 at 4 µM was incubated in the presence of 10 µM of poly-dT50 in MN buffer containing 1 mM ATP with 10 mM $MgCl_2$ and imaged. No droplets and no fluorescence signal could be detected.

**C.** Microscopy analysis of *Vc*DciA-ATTO 647 mixed with *Vc*DnaB-ATTO 550. *Vc*DciA-ATTO 647 at 15 µM was mixed with *Vc*DnaB-ATTO 550 at 4 µM in presence of 10 µM of poly-dT50 in MN buffer containing 1 mM ATP with 10 mM $MgCl_2$. The mixed solution was loaded on the Pluronic pretreated glass and imaged.

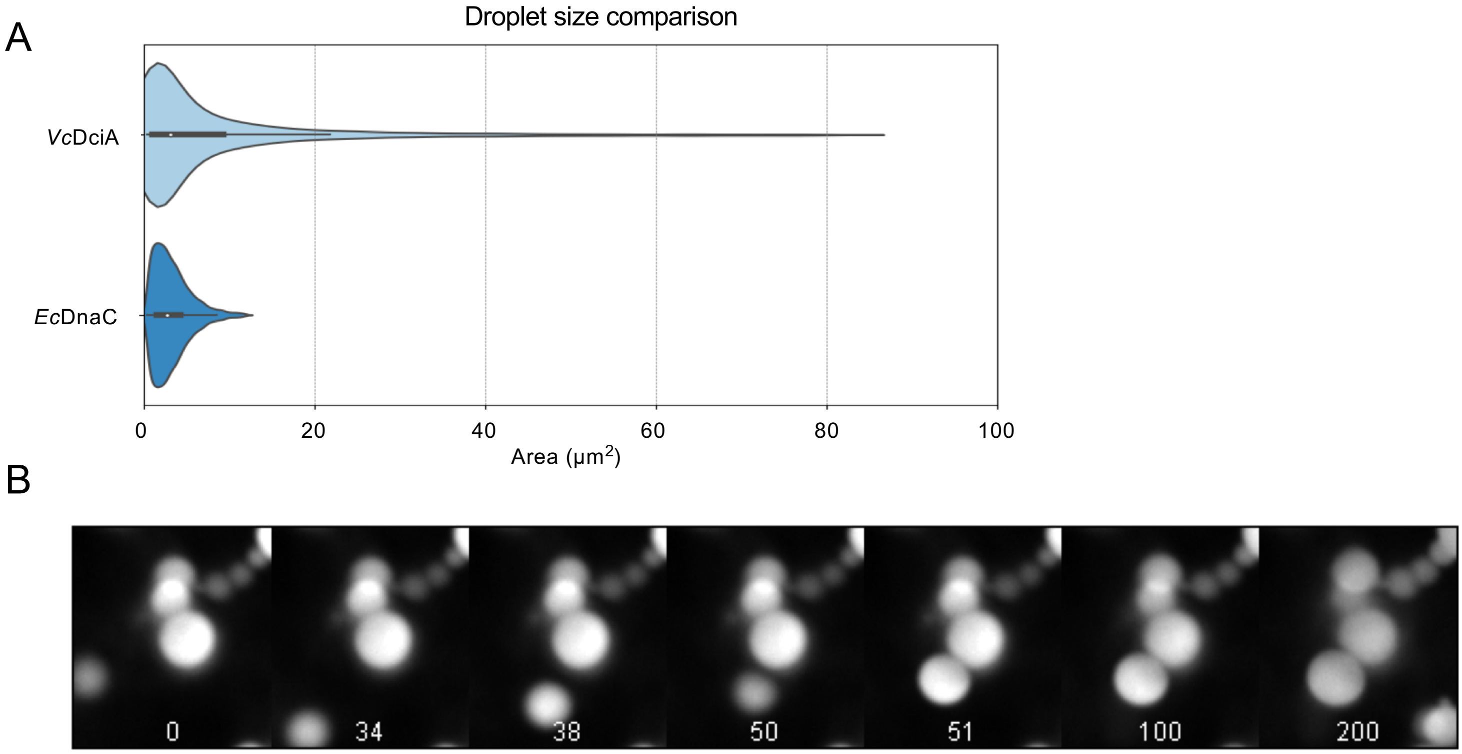


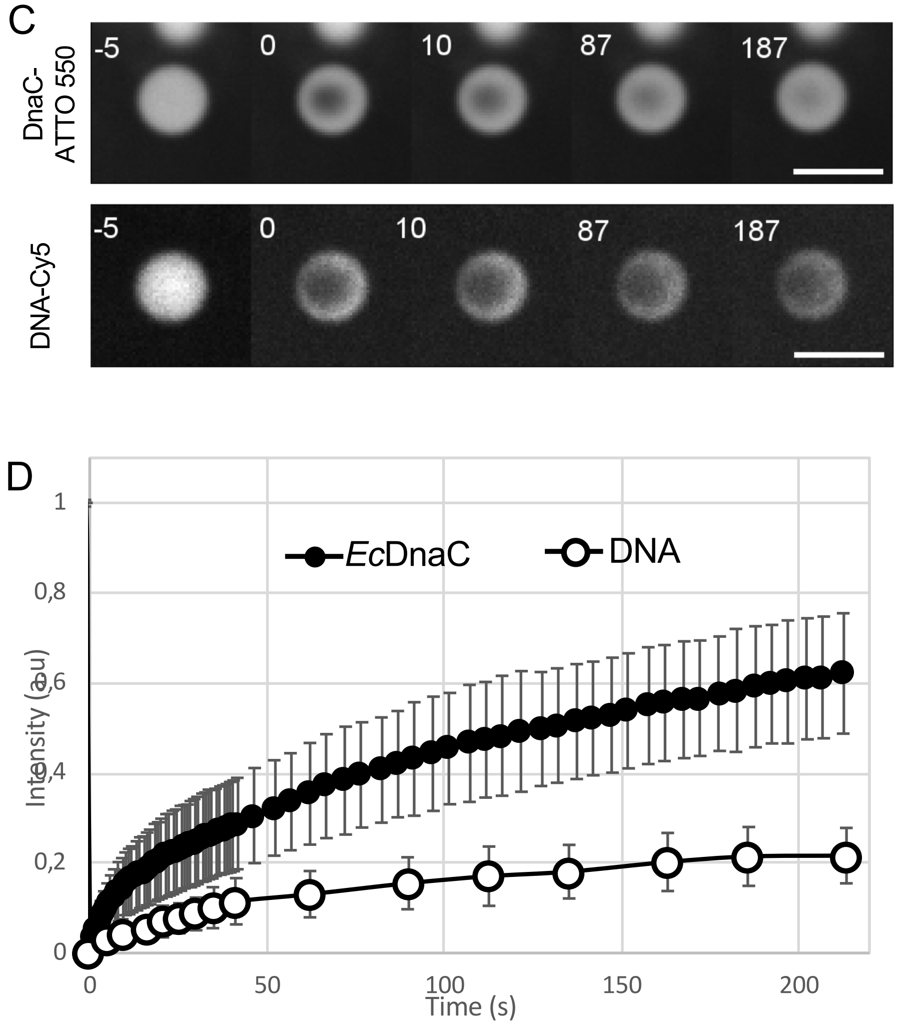


**Supplementary Figure 4 : *Ec*DnaC exhibits a higher mobility than DNA within a droplet, such as *Vc*DciA.**
LLPS were first formed with 30 µM of *Ec*DnaC-ATTO 550 in the presence of 10 µM of poly-dT50 and 0.2 µM of ssDNA-Cy5.
**A.** Size comparison. Violon plots that represent *Vc*DciA sizes (n= 143) and *Ec*DnaC sizes (n= 78). The area is in µm$^2$
**B.** *Ec*DnaC droplets do not fuse. Example of two droplets touching each other without fusing or two droplets fusing but not relaxing into one unique bigger droplet.
**C.** Time-lapse imaging of the same *Ec*DnaC droplet after FRAP experiments. FRAP was applied on a central sub-region (1 µm diameter). Top raw is the tagged protein and bottom raw is the tagged ssDNA. Time is indicated in seconds. The bleaching event is performed at the same time for both channels: time 0s. The scale bar is 3 µm.
**D.** Quantification of relative intensity recovered after FRAP experiments. Intensity recoveries of the mean fluorescence (± Standard deviation) of *Ec*DnaC-ATTO 550 (full circles) and DNA-Cy5 (open circles) are presented. n=34 on four independent experiments.
